# Females and hermaphrodites of the gynodioecious *Geranium maculatum* respond similarly to soil nutrient availability

**DOI:** 10.1101/2020.08.21.261305

**Authors:** Katharine Putney, Mavis Wolf, Chase Mason, Shu-Mei Chang

**Affiliations:** Department of Biology, Wofford College, 429 N Church St, Spartanburg, SC 29303; Department of Plant Biology, University of Georgia, 120 Carlton Street, Athens, GA 30602; Department of Biology, University of Central Florida, 4110 Libra Drive, Orlando, FL 32816

**Keywords:** Geranium, gynodioecy, nitrogen, nutrients, phosphorus

## Abstract

Sexual dimorphism in plant growth and/or reproductive responses to the surrounding environment has been documented in some plant species. In gynodioecious plants, it is especially important to understand whether females and hermaphrodites differ in their response to environmental stressors, as the fitness of females relative to hermaphrodites determines the extent to which these separate sexes are maintained in natural populations. Soil nutrient availability is of particular importance given the different nutrient requirements of male and female sexual functions in plants. Here, we evaluated and compared the growth of females and hermaphrodites of *Geranium maculatum* in response to varying levels of nutrients. Using a greenhouse experiment, we manipulated the overall nutrient, nitrogen, and phosphorus levels in the soil and measured growth, allocation, and leaf quality responses in both females and hermaphrodites. We found that sexes responded similarly in their growth and allocation responses to nutrient availability, albeit evidence that female leaf chlorophyll content may have increased more than that of hermaphrodites across soil nitrogen levels. Our findings demonstrate that hermaphrodites differ from females in terms of their physiological response to varying nutrient levels, however these physiological differences did not translate into meaningful growth or reproduction differences.

## Introduction

Sexual dimorphism is common and often conspicuous in the animal kingdom (Hedrick and Temeles, 1989), which is often attributed to sexual selection processes that favor divergence in traits important for mate attraction. For dioecious plant species, those with male and female individuals, similar sexual selection processes can also lead to sexual dimorphism in floral traits. Additionally, fertility selection, in which individuals may employ an optimal allocation strategy to maximize their fitness gain, could lead to morphological differences between sexes (Charnov et al., 1976; Delph, 1999; Delph and Herlihy, 2012). Sexual dimorphism in plants, however, is expected to be more subtle since sexual selection may be constrained by the limited ways in which female choice can be exhibited in plants (Skogsmyr and Lankinen, 2002). Additionally, the enhanced competition due to common limiting resources between sexes, such as light, resources, and pollinator services, may restrain the effect of divergent selection and reduce the variation between sexes in certain instances (Labouche and Pannell, 2016). Despite this, examples of dimorphism in plants are still found for reproductive traits (Eckhart, 1999), vegetative traits (Barrett and Hough, 2013), physiology (Dawson and Geber, 1999), and even in growth responses to environmental changes (Petry et al., 2016).

Sexual dimorphism in plant growth and/or reproductive response to changes in the environment has serious ecological and evolutionary implications, especially if these growth differences lead to differences in survival rate or reproductive output under certain environmental conditions. In dioecious plants, populations may decline if one sex is severely underrepresented and becomes a limiting factor for reproductive success of the other sex (Le Galliard et al., 2005; Shelton, 2008). Dimorphism in growth response can also affect the evolutionary trajectory of other sexual systems, such as gynodioecy. Gynodioecious plant species have hermaphroditic as well as male-sterile female individuals in some or all populations. Gynodioecy is often considered an evolutionary “stepping-stone” between hermaphroditism and dioecy (Barrett, 2002). The maintenance of gynodioecy and other breeding systems is likely facilitated by ecological agents of selection, such as water and nutrient availability, as hermaphrodites may optimize lifetime fitness by allocating more resources to pollen production than seed production during periods of environmental stress (Sex-Differential Plasticity hypothesis, e.g., see Delph, 2003). While hermaphrodites may exhibit this type of plasticity in resource allocation, females are expected to maintain relatively more constant seed production across environments as seed production is the sole avenue to fitness in female plants. This creates a situation in which the seed production threshold females must achieve in order to compensate for the lack of male fitness would be lower in resource poor habitats than in resource rich habitats, thus explaining why females in gynodioecious systems are often found in harsh environments (Delph, 2003). Testing the extent to which females and hermaphrodites differ in their responses to environmental stresses is, thus, critical in understanding how and whether females can be maintained across generations within a gynodioecious species.

While both hermaphrodites and females produce fruits and seeds, they differ inherently in their ability to produce pollen, which has higher nitrogen and phosphorus requirements than fruit and seed production per unit mass (e.g. Ishida et al., 2005). In addition, fruiting structures often have some photosynthetic capability to support seed production (e.g. Galen et al., 1993) but pollen-bearing structures are rarely photosynthetic. Hence, male function unique to hermaphrodites may result in differential growth responses to nutrient limitation, due to the direct cost of pollen production and perhaps also due to additional costs of the larger flowers and floral displays typical of hermaphrodites relative to females (Delph, 1996). If we assume that any trade-offs with other physiological functions (WUE, photosynthetic capacity, etc) will ultimately affect growth via the efficiency of carbon assimilation, this overall cost of male function could have the potential to reduce other growth investments, such as vegetative growth or female function. However, such trade-offs are not a general rule, and several previous studies, including one on our study species, either did not find significant differences between the sexes in photosynthetic rate or water use efficiency, or those differences were very small (Caruso et al., 2003; Varga and Kytöviita, 2017). Given a lack of physiological differences and, hence, the ability to assimilate carbon, one would expect that hermaphrodite growth and/or reproductive performance would be more sensitive to lower nutrient levels than that of females (Eckhart and Chapin, 1997).

Few studies have directly evaluated sexual dimorphism in response to nutrient availability in gynodioecious plants, but the available data indicate either that hermaphrodites tend to show a stronger response to nutrient limitation than females, or that there is little to no difference in response. In a greenhouse experiment on the annual gynodioecious species *Phacelia linearis*, hermaphrodites allocated a greater proportion of biomass to belowground structures than did females, and this difference was significantly greater under low than high nitrogen availability (Eckhart and Chapin, 1997). As higher allocation to belowground growth is an expected growth response to low nutrient availability (Ericsson, 1995; Hunt and Lloyd, 2008), these findings suggest that hermaphrodites in this species were likely experiencing more nutrient limitation at low nutrient levels than were females. Additionally, in the perennial herb *Plantago lanceolata*, when females and hermaphrodites were grown under nitrogen-limiting conditions, females had higher aboveground dry mass than hermaphrodites, despite similar nitrogen uptake (Poot, 1997). This is also consistent with a growth cost of male function, as a portion of the nitrogen taken up by hermaphrodites was used for pollen production rather than put towards photosynthetic biomass.

In this study, we used *Geranium maculatum* as our model system. To our knowledge, there are no other studies that directly evaluate the sexes responses to manipulations of separate components of fertilizer in this species, however there are a handful of relevant studies in closely related species. In *Geranium sylvaticum*, for example, Asikainen & Mutikainen (2005) found that increased overall soil nutrients resulted in increased seed production for both sexes equally. In addition, Varga et al. (2008) found evidence for sex-specific interactions with mycorrhizal fungi, mutualists that are a critical component of plant nutrient relations in the field. More specifically, they found that females produced more seeds of equal phosphorus content (per seed) than hermaphrodites, and that seed production was positively correlated with the number of fungal structures in the roots most closely linked to phosphorus exchange (arbuscules). In another closely related species, *G*. *richardsonii*, Williams et al (2000) found that flower size was consistently different between the two sex morphs, despite the effects of available sunlight. Shaded plants had larger flowers, regardless of the sex, and hermaphrodites still had larger flowers in the shade.

Given the potential for differences between the sexes in their relationship to nutrient availability, in what ways do females and hermaphrodites differ in their growth and biomass allocation responses to changes in nutrient availability? We addressed this question by evaluating the growth of females and hermaphrodites of *Geranium maculatum*, a perennial gynodioecious plant species, across various levels of nutrients. We predicted that as nutrient availability decreases, both sexes would decrease in biomass, increase allocation to belowground growth, and the quality of their leaves in terms of nutrient concentrations would decrease. In addition, we predicted that hermaphrodite biomass, allocation, and leaf quality would change to a greater extent than those traits in females due to the cost of male function.

## Materials and Methods

### Study System

*Geranium maculatum* is a perennial rhizomatous herb that grows throughout the eastern half of North America ranging from Ontario, Canada to Alabama and Georgia, and west to Missouri (Weakley et al., 2013). Similar to other gynodioecious species, some *G*. *maculatum* populations consist only of hermaphrodites (which produce both ovules and pollen), while other populations contain a mix of both hermaphrodites and females (male-sterile individuals) that solely produce ovules (Ågren and Willson, 1991).

In this experiment, we used plants from three sexually dimorphic populations from Southeastern US: “Azalea Road” (AZ) from Asheville, North Carolina (latitude: 35.574268, longitude: -82.489967), “Bear Hollow” (BH) in Athens, Georgia (latitude: 33.926727, longitude: -83.386391), and “Ellijay” (EL) in Ellijay, Georgia (latitude: 34.770084, longitude: -84.589966).

### Plant preparation

Plants used in this study came from a larger collection curated by our group and had been growing in common conditions for multiple years, thus decreasing the environmental effects linked to their home site, such as differential stored nutrients or sugars due to differential sunlight or nutrient availability. This method would not be able to eliminate genetic differences among populations, however, so population origin was still factored into the statistical analysis (see “Statistical Analysis” below). By cold stratifying rhizomes at 5°C for roughly one month following approximately four months of growth inside the greenhouse, we were able to simulate a new season’s growth for our experimental plants. We exposed 92 genotypes to cold stratification (13 females and 14 hermaphrodites from AZ; 18 females and 21 hermaphrodites from BH; and 12 females and 14 hermaphrodites from EJ). Afterward, rhizomes with more than one active meristem for new ramets were split as evenly as possible to produce clones for this experiment. Most genotypes were represented by no more than three clones, producing a total of 175 experimental plants (19 females and 23 hermaphrodites from AZ; 43 females and 47 hermaphrodites from BH; and 19 females and 24 hermaphrodites from EJ). After cloning, excess dirt was removed before rhizomes were weighed with roots attached (initial belowground fresh mass). Following these steps, we found that the initial belowground fresh mass was similar between the two sexes (F_1,166_=1.82, p=0.18).

Because all experimental plants had been previously exposed to outdoor environments in Athens, GA for multiple seasons, in order to isolate the nutrient response of the plant it was necessary to remove roots from rhizomes to eliminate any possible bacterial or fungal symbionts present that could affect plants’ nutrient acquisition. We removed roots just prior to planting. Rhizomes were then rinsed in clean water to remove excess dirt, dipped briefly in a 3% bleach solution to kill any remaining bacteria/fungal spores on the rhizome surface, and then immediately rinsed off again in clean water to remove the bleach. This process has been demonstrated to result in *G*. *maculatum* plants with no arbuscular mycorrhizal fungi in their roots once new roots have sprouted, although a very small amount of septate fungi (non-AM fungi) may remain (Quigley, 2013). The cleaned rhizomes were then each shallowly planted in 1.67-liter pots (15.24 cm diameter, 14.6 cm in height; ITML 6-inch Regal standard pots, BFG supply company, Burton, OH, USA) with approximately one liter of steam-sterilized sand. Pots were placed in separate 10-cm deep saucers (25.4 cm in diameter) and watered to field capacity. Plants were exposed to ambient light in the greenhouse at approximately 24 C daytime and 18 C nighttime temperatures.

### Nutrient treatments

In order to control overall nutrient content and independently manipulate nitrogen and phosphorus, we created our own nutrient solutions based on a half-strength standard Hoagland’s recipe. The original recipe includes appropriate amounts of both macro-and micro-nutrients required for normal plant growth (Hoagland and Arnon, 1950). We modified either the nitrogen content or phosphorus content while keeping all other nutrients constant by manipulating the nitrate/ammonia and phosphorus containing compounds, respectively (“nitrogen or phosphorus treatments” hereafter). Nitrogen was manipulated such that the ratio of nitrate to ammonia was kept at a constant 7 : 1 ratio in all manipulations. We made Hoagland’s solutions with the following nitrogen concentrations in mmol/L: 0.4, 2, 8 (control), and 16. We also made solutions with the following phosphorus concentrations: 0.01, 0.5, 1, and 4 (Fig. 1, Supp. Table 1), with 1 representing the control, same solution as the 8 mmol/L N treatment. In addition to these varying nitrogen and phosphorus concentrations, we also created three dilution treatments of 1X (half-strength Hoagland’s), 0.5X, and 0.1X by diluting the half-strength modified Hoagland’s solution (1X) for each of the nitrogen or phosphorus treatment solutions. As such, we created a factorial design of primary nutrient concentration and overall Hoagland’s solution dilution for each focal nutrient (N and P; Fig.1).

**Figure 1.**
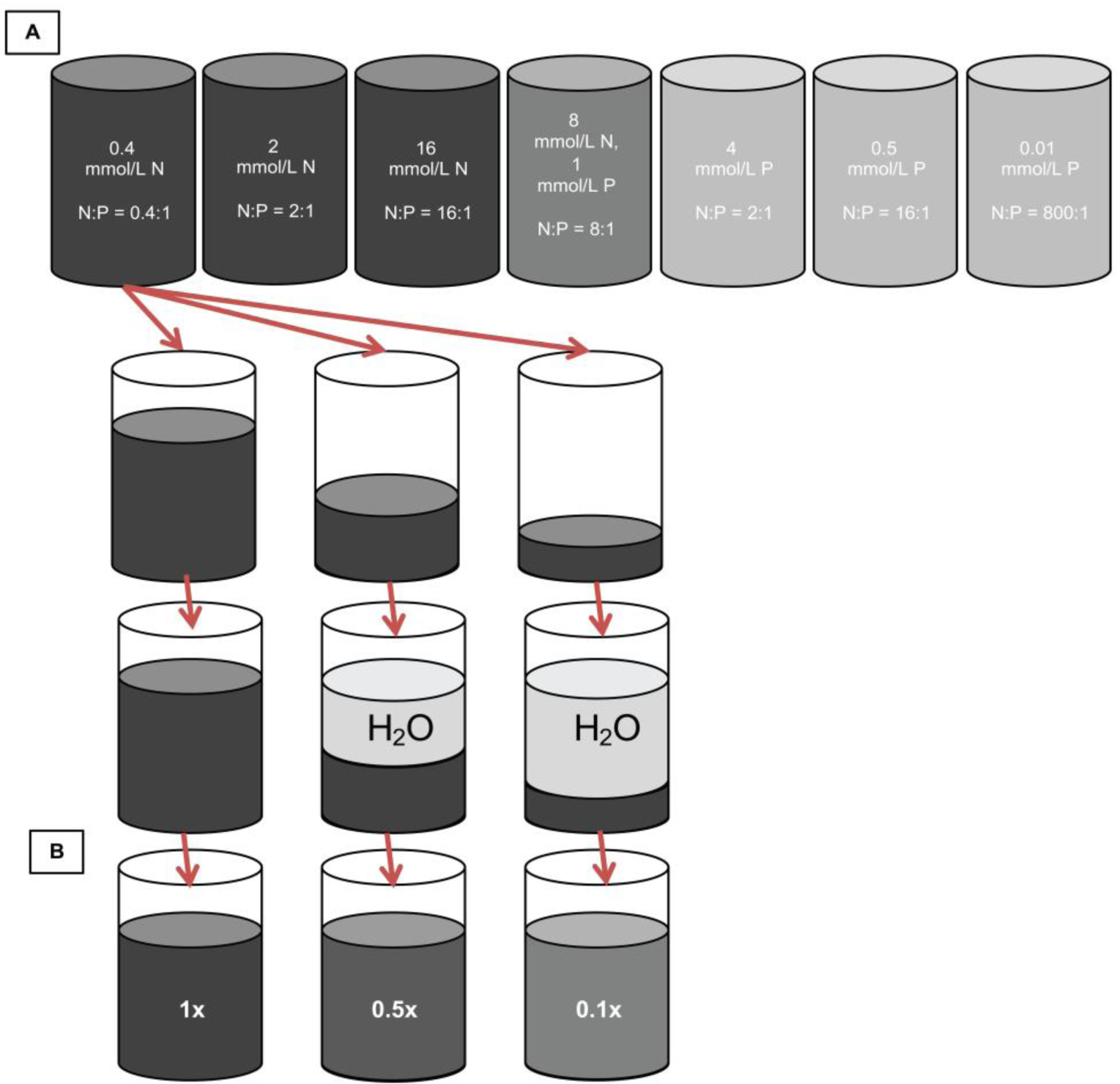
Diagram of the creation of treatments. Starting from a ½ strength Hoagland’s recipe with an N:P ratio of 8:1 (green vial), we further modified the recipe’s N:P ratio by changing either nitrogen (blue vials) or phosphorus (purple vials), for a total of 7 N:P ratio solutions (A). Each of these N:P ratio solutions was then separated into three separate solutions which were either not diluted (1x), diluted to half the original strength (0.5x) or diluted to one-tenth the original strength (0.1x) (B).

To ensure non-limiting water availability during the course of the growing period, each plant was watered and fertilized using the following method. Before the initial treatment application, all plants were watered to field capacity and excess water was removed from the saucers. Each plant was subsequently watered with 500mL of fertilizer solution; 250mL added directly into the pot (achieving saturation), and the remaining 250mL poured into the saucer. We kept the amount poured into the saucer consistent to keep any effects of evaporation of that portion of the fertilizer consistent across plants. Over the course of the growing period, plants were fertilized with the assigned treatment solution approximately once a month. To prevent the plants from drying out between treatment applications, extra watering was applied as needed (to all plants each time) at a constant volume of 500mL per plant, split similarly as the previously described fertilizer solutions. Need for watering was determined by whether at least one plant had just begun to visually lose turgor in the leaves and occurred approximately once a week for weeks when we were not applying the fertilization treatment. We used a complete randomized block design such that plants within each treatment combination were represented in each block and their spatial position was randomized within a given block.

### Data Collection

Over the course of approximately nine and a half weeks, we took weekly measurements of leaf count, flower count, and senesced tissues. SPAD values of green leaves (a proxy for relative leaf chlorophyll content; (Minolta, 1989) were measured at the end of the nine and a half weeks using a SPAD 502 Plus Chlorophyll Meter (Spectrum Technologies, Inc.). A mean SPAD value for each plant was determined by taking three measurements per leaf on a maximum of three fully expanded but not-yet senesced leaves and averaging these values for a given plant (Richardson et al., 2002). For plants with less than three green leaves, we took three measurements per leaf for all leaves possible and averaged these values for that plant.

At the end of the nine and half weeks, all aboveground biomass was harvested. For each individual plant, we removed all aboveground structures while leaving the rhizome intact in the pot. Aboveground tissues were separated into the following categories: whole inflorescence, green leaves, senesced leaves, and petioles. Whole inflorescence mass included flowering stems (any stems with branching or containing more than one leaf) and all live and senesced flowers. Green leaves were separated from their respective petioles, weighed for fresh mass for leaf dry matter content calculations, and then scanned using an Epson Perfection 3490 PHOTO scanner. Total green leaf adaxial area was calculated using ImageJ (Schneider et al., 2012). All three tissue types were then individually dried at 60°C for approximately four days and then weighed. After green leaf tissue was weighed, one portion was analysed for percent phosphorus with colorimetry (Varvel et al., 1976), while another portion was analysed for percent nitrogen using Micro-Dumas combustion by the Stable Isotope Lab at the University of Georgia (http://siel.uga.edu/).

To obtain belowground biomass, potted rhizomes and their associated roots were first excavated, rinsed in water to remove excess sand, air-dried for approximately 20 minutes, and then weighed for fresh mass. In order to get values more comparable to other plant parts for questions of allocation, a small portion of rhizome was extracted from one typical rhizome which was weighed both before and after drying at 60°C for one week. We then used the proportional change in mass from this sample to adjust belowground fresh mass to a dry mass estimate. We used the fresh rhizome mass for any analyses that involved only belowground biomass but used the estimated rhizome dry mass when it is combined with other structures (such as total biomass or relative allocation of above and below ground mass) in order to make it comparable to the dry mass measurements for other organs.

At the time of plant collection, 100% of plants still had a live rhizome that could be weighed, however only 83% of plants had sprouted to produce leaves, and 42% had flowered. Aboveground growth was just beginning to senesce on a majority of plants at this time, so we ended the experiment in order to get the most accurate measurement of aboveground biomass. It is also worth noting that stem and leaf growth in the field usually takes about 30-40 days to reach maximum growth, which is a similar time period we allowed for our plants (Martin, 1965).

### Statistical analysis

All analyses included the source population of the plant, plant sex (Female or Hermaphrodite), dilution (1x, 0.5x, 0.1x), and nutrient treatment (0.4N, 2N, 8N, and 16N for nitrogen analyses; 0.01P, 0.5P, 1P, and 4P for phosphorus analyses; all seven nutrient treatments for dilution analyses) as predictor variables, and initial rhizome mass as a covariate. To evaluate whether the sexes differed in their response to dilution or nutrient treatments, we also included the interaction term of sex and either factor as additional predictors. We also included a dilution and nutrient treatment interaction to account for the possibility that dilution effects depend on the relative N or P, and vice versa. We did not include the three-way interaction of sex, dilution, and nutrient treatment due to insufficient sample size. In addition, block was dropped from the final analyses because preliminary results showed that its effects were consistently non-significant. Due to the use of cloning, plant genotype was considered for the analysis. However, clones of genotypes were distributed across treatments such that there was no replication of any given genotype in any given treatment or treatment combination, so we did not have the power to detect its effects in our analysis. All factors were treated as fixed variables (except block, prior to being removed from the analysis). To address questions concerning the effect of nutrient dilution, the entire dataset was used in the analysis. In contrast, for questions specific to nitrogen or phosphorus manipulation, we analyzed the subset of data for which that specific nutrient had been manipulated, plus the 8 mmol/L N, 1 mmol/L P control. (See Supplemental Table 2 for models and datasets used for each trait and question).

We tested for normality of residuals produced by univariate analysis of the raw data for each trait using a Shapiro-Wilk test. For traits that violated the normality assumption of their residuals, we used the transformed data in the final analyses. All residuals from the analyses using transformed data satisfied the normality assumptions. Specifically, we analyzed the square root transformation of leaf number and flower number, the arc sin square root transformation of percent leaf phosphorus and the ratio of inflorescence to total dry mass, and the log transformation of leaf N:P ratio. Due to the large number of tests within our study, we determined P-value significance using Benjamini-Hochberg adjustments at a false discovery rate of 0.10 (Benjamini and Hochberg, 1995).

## Results

### Biomass

Consistent with expectations, overall soil nutrient level (i.e. dilution treatment) had an effect on the total biomass of both sexes in *G*. *maculatum*. Total plant dry mass was significantly lower in the 0.1x treatment than in the 1x and 0.5x treatments (total dry mass: F_2,138_=4.85, P=0.0092; Fig. 2a). This pattern was driven primarily by aboveground dry mass (F_2,109_=8.21, P=0.0005; Fig. 2b), as belowground fresh mass was not significantly different among dilution treatments (Fig. 2c). The sexes did not differ significantly in their biomass across dilution treatments (F_2,138_=1.96, P=1.45). Aboveground dry mass of both sexes also decreased as relative soil nitrogen concentrations decreased (F_3,62_=5.42, P=0.0022; Fig. 2e), whereas biomass did not change in response to relative soil phosphorus concentration (F_3,76_=0.20, P=0.89). Again, the sexes did not differ in their biomass across soil nitrogen (F_3,79_=0.92, P=0.43) or soil phosphorus levels (F_3,75_=2.13, P=0.10).

**Figure 2.**
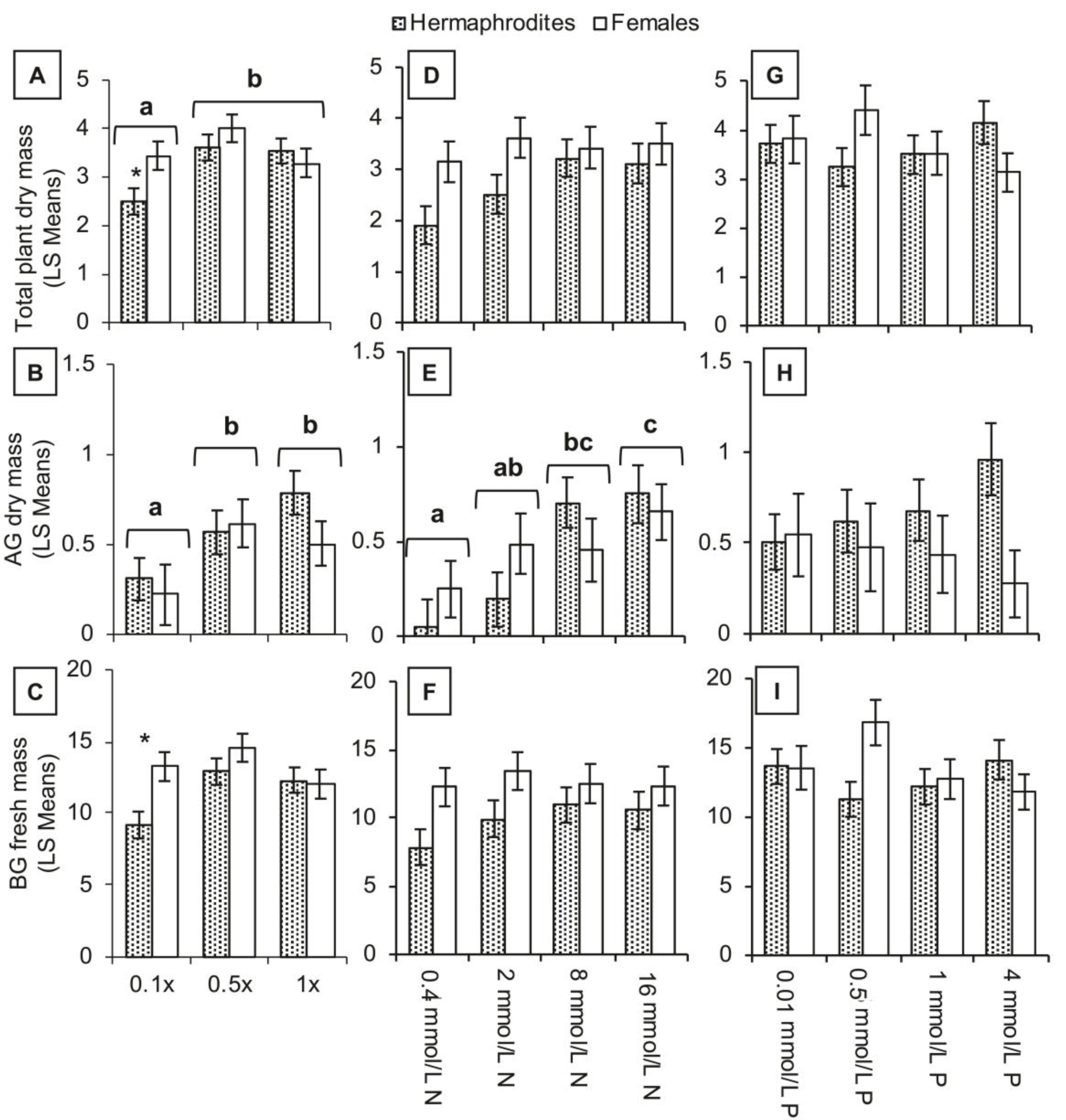
Least squares means of total plant dry mass (g; A, D, G), aboveground dry mass (g; B, E, H), and belowground fresh mass (g; C, F, I) of the two sexes across the three dilution treatments (A, B, C), the four nitrogen treatments (D, E, F), and the four phosphorus treatments (G, H, I). Pairs of bars with different letters within a trait are significantly different from each other. Pairs of means with asterisks above are those in which females and hermaphrodites are significantly different from post-hoc comparisons of means.

### Reproduction

Inflorescence dry mass of both sexes also decreased as overall soil nutrients decreased (F_2,105_=4.37, P=0.015; Fig. 3a), however flower number did not significantly change as overall soil nutrient levels decreased (F_2,138_=1.96, P=0.14; Fig. 3b). In contrast, both inflorescence dry mass and flower number decreased as relative soil nitrogen concentration decreased (infl. dry mass: F_3,58_=3.34, P=0.025; flower number: F_3,79_=3.17, P=0.029; Fig. 3c and d). These reproductive measures did not change across relative soil phosphorus concentrations (infl. dry mass: F_3,59_=0.19, P=0.91; flower number: F_3,75_=0.93, P=0.43). The sexes did not differ significantly in their reproductive responses to varying levels of overall soil nutrients (infl dry mass: F_2,105_=0.27, P=0.76; flower number: F_2,138_=1.14, P=0.32), soil nitrogen (infl dry mass: F_3,58_=2.24, P=0.09; flower number: F_3,79_=0.48, P=0.70), or soil phosphorus (infl dry mass: F_3,59_=0.41, P=0.28; flower number: F_3,75_=1.28, P=0.29).

**Figure 3.**
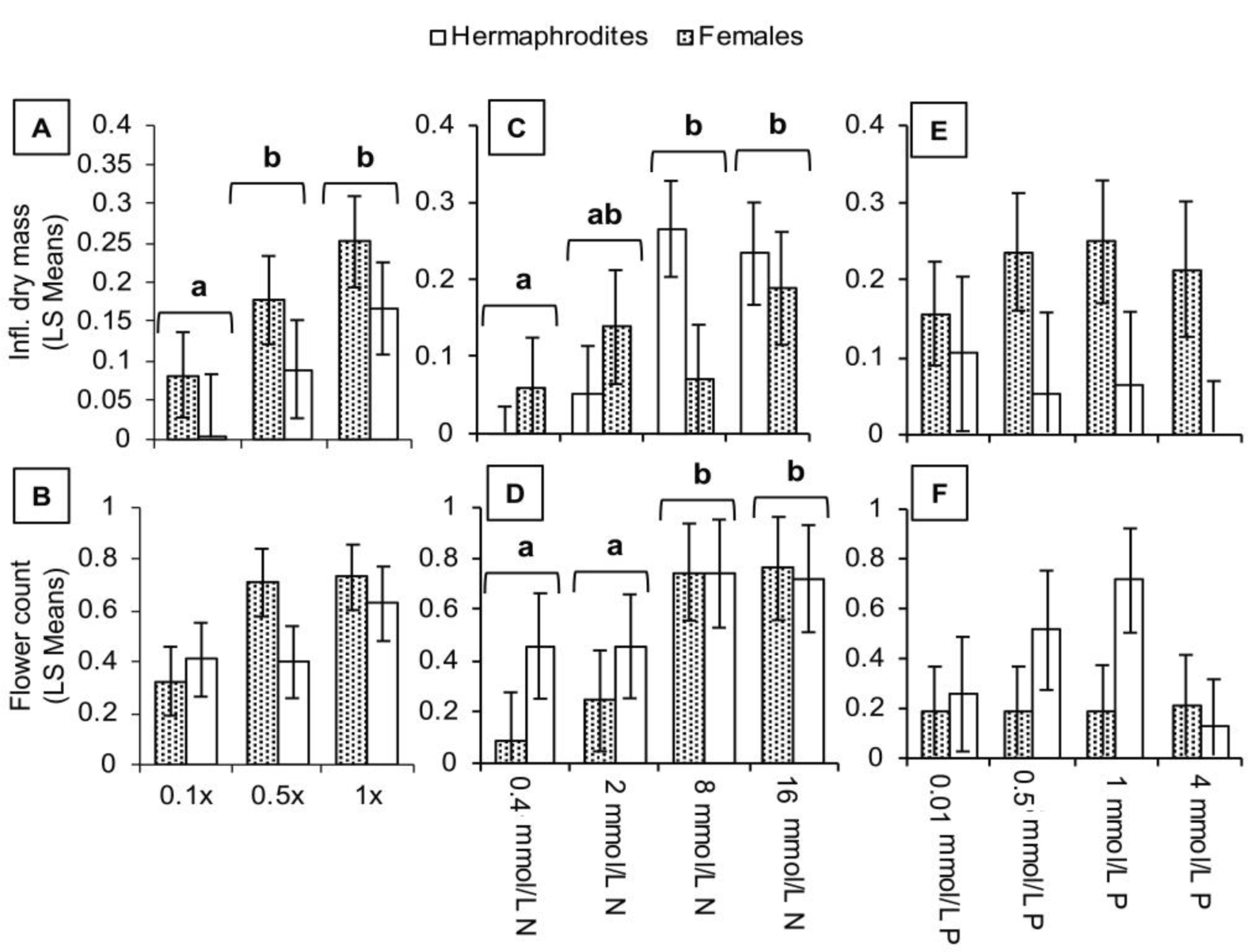
Least squares means of inflorescence dry mass (g; A, C, E) and highest recorded flower count at one time (B, D, F) of the two sexes across the three dilution treatments (A, B), the four nitrogen treatments (C, D), and the four phosphorus treatments (E, F). Pairs of bars with different letters within a trait are significantly different from each other. Pairs of means with asterisks above are those in which females and hermaphrodites are significantly different from post-hoc comparisons of means.

### Biomass allocation

Relative allocation to belowground biomass decreased as overall soil nutrient level increased for both sexes (F_2,138_=3.84, P=0.024; Fig. 4a) and relative allocation to inflorescences likewise decreased as soil nutrients decreased (F_2,138_=5.68, P=0.0043; Fig. 4b). In response to relative soil nitrogen concentration, however, only relative allocation to belowground biomass changed (F_3,79_=4.63, P=0.0049). As soil nitrogen decreased, relative allocation to belowground biomass increased (Fig. 4c). The sexes did not differ in their allocation responses to overall soil nutrient levels (BG allocation: F_2,138_=1.94, P=0.15; infl allocation: F_2,138_=0.33, P=0.72) and soil nitrogen levels (BG allocation: F_3,79_=0.34, P=0.80; infl allocation: F_3,79_=2.22, P=0.09), and neither of the sexes’ allocation patterns changed in response to relative soil phosphorus levels (main effect: BG allocation: F_3,75_=0.95, P=0.42; infl allocation: F_3,75_=1.22, P=0.31; sex*phosphorus: BG allocation: F_3,75_=1.59, P=0.14; infl allocation: F_3,75_=1.27, P=0.58).

**Figure 4.**
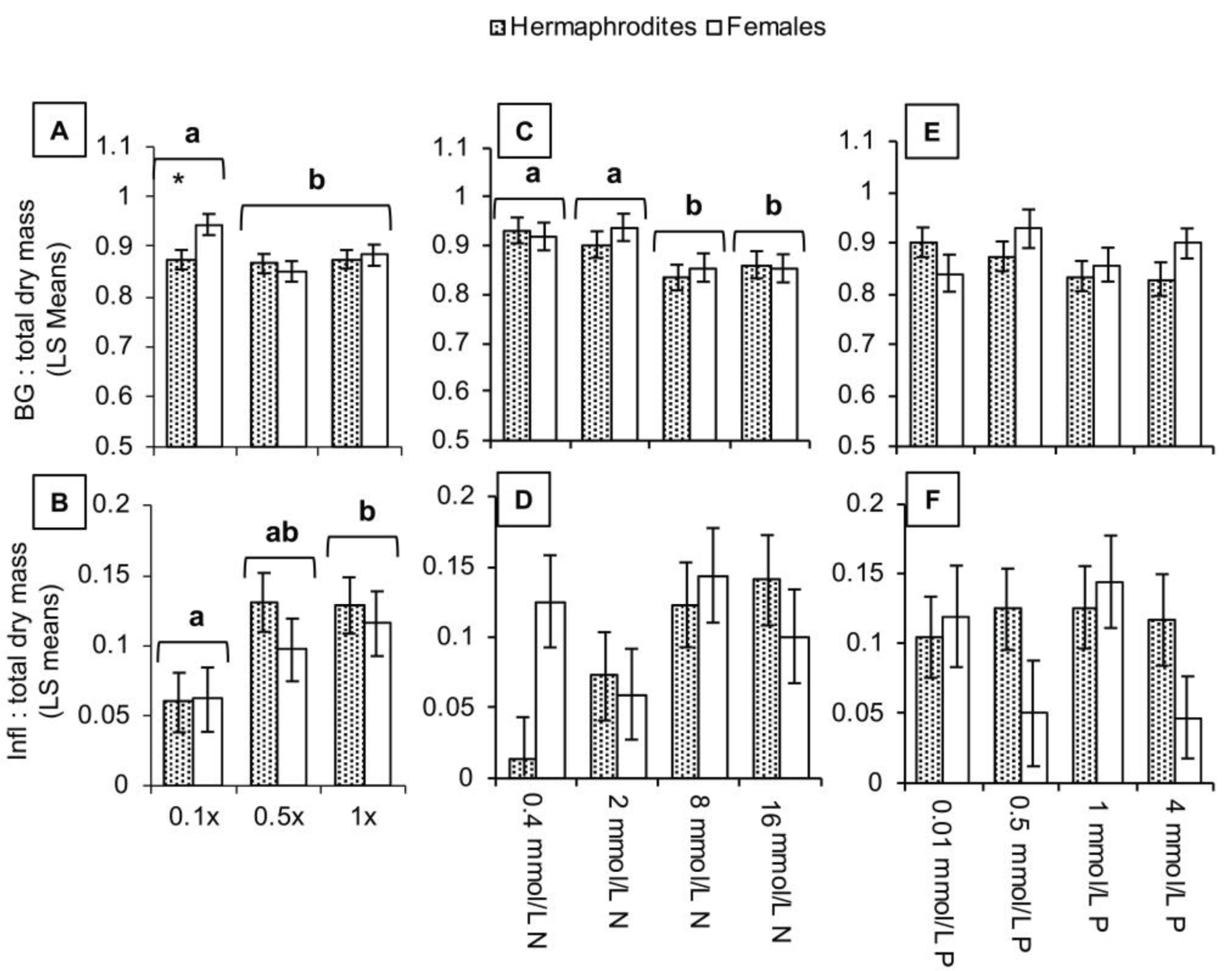
Least squares means of the ratio of belowground to total plant dry mass (A, C, E) and the ratio of inflorescence to total plant dry mass (B, D, F) of the two sexes across the three dilution treatments (A, B), the four nitrogen treatments (C, D), and the four phosphorus treatments (E, F). Pairs of bars with different letters within a trait are significantly different from each other. Pairs of means with asterisks above are those in which females and hermaphrodites are significantly different from post-hoc comparisons of means.

### Leaf quality

Leaf nitrogen and SPAD values responded to treatments, whereas all other leaf quality traits did not. Leaf nitrogen decreased as overall soil nutrients decreased (F_2,64_=5.07, P=0.0090; Fig. 5b), and as relative soil nitrogen levels decreased (F_3,37_=4.84, P=0.0061; Fig. 5f). SPAD values also decreased as relative soil nitrogen decreased (F_3,46_=4.08, P=0.012), however the SPAD values of the two sexes responded differently to soil nitrogen. Specifically, female SPAD values were highest at the 8mmol/L N treatment, whereas hermaphrodite SPAD values did not vary significantly across nitrogen levels (F_3,46_=5.81, P=0.0019; Fig. 5h). Leaf nitrogen and SPAD values were not different across relative soil phosphorus levels (main effect: Leaf N: F_3,31_=1.98, P=0.14; SPAD: F_3,39_=0.97, P=0.42; sex*phosphorus: Leaf N: F_3,31_=2.15, P=0.11; SPAD: F_3,39_=1.12, P=0.35). Complete reporting of statistical results can be found in Supplemental Table 2a-c.

**Figure 5.**
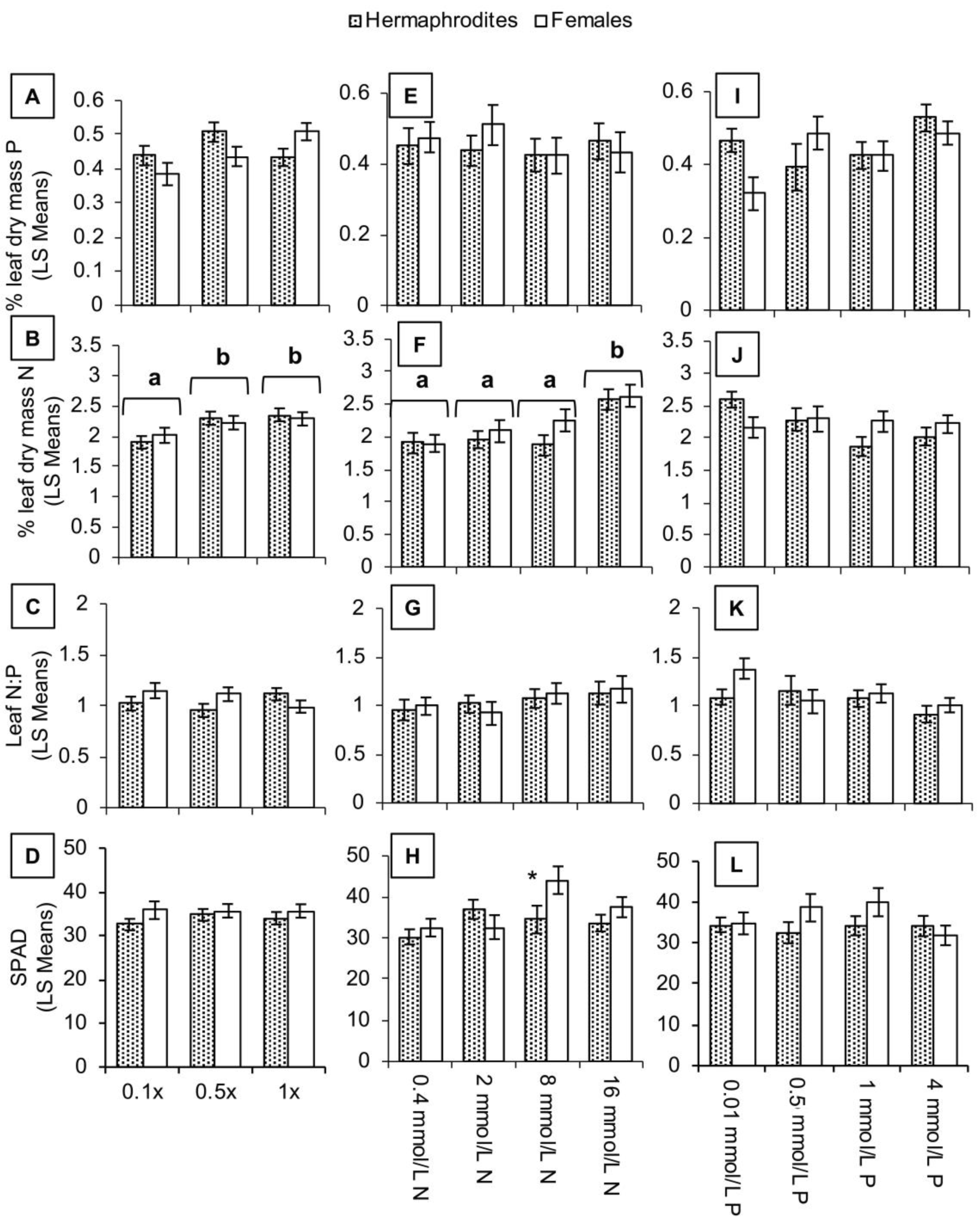
Least squares means of per cent phosphorus by leaf dry mass (A, E, I), per cent nitrogen by leaf dry mass (B, F, J), the ratio of per cent leaf nitrogen to leaf phosphorus (C, G, K), and leaf SPAD values (D, H, L) of the two sexes across the three dilution treatments (A, B, C, D), the four nitrogen treatments (E, F, G, H), and the four phosphorus treatments (I, J, K, L). Pairs of bars with different letters within a trait are significantly different from each other. Pairs of means with asterisks above are those in which females and hermaphrodites are significantly different from post-hoc comparisons of means.

**Figure 6.**
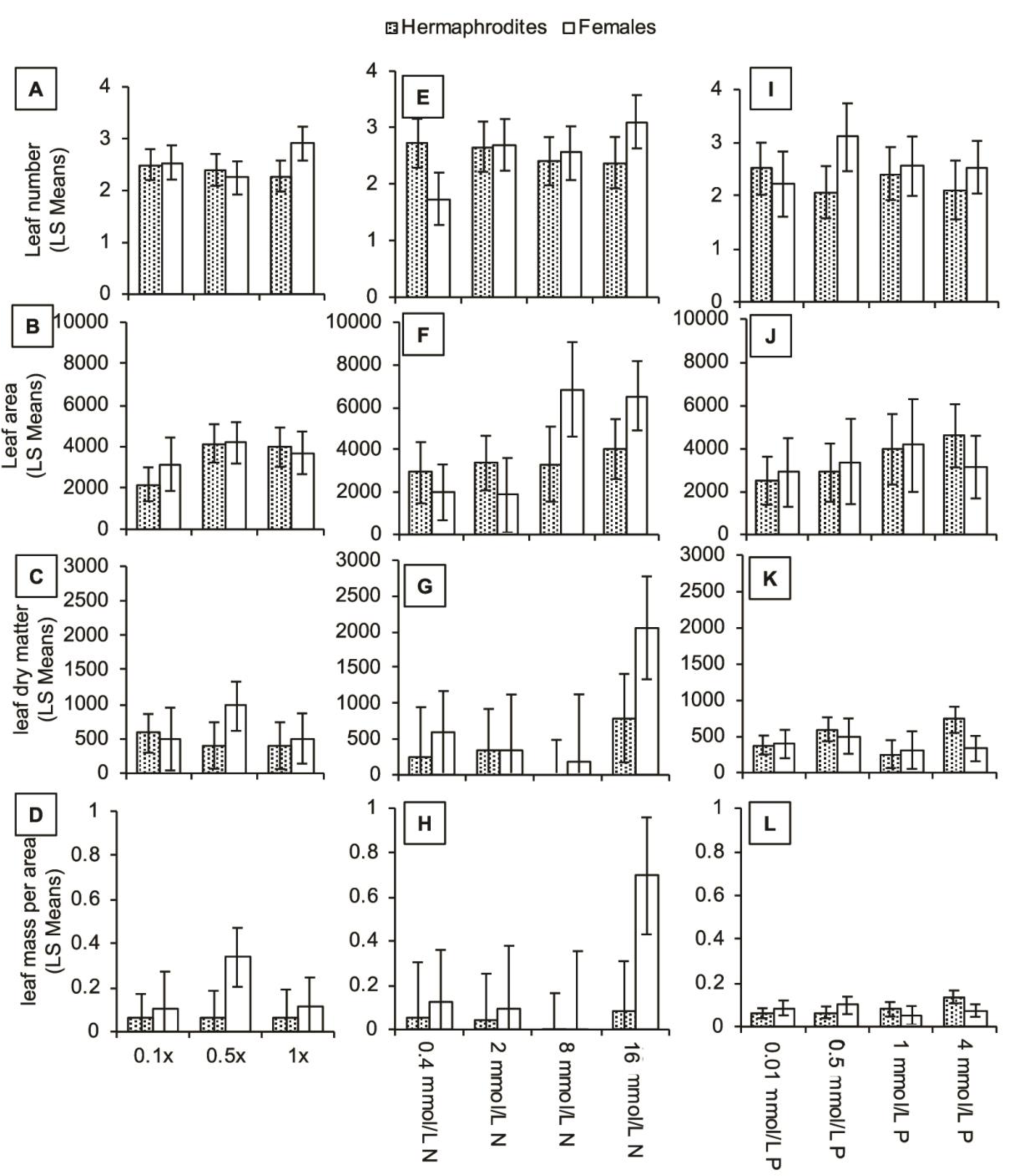
Least squares means of the highest recorded leaf number at one time (A, E, I), total green leaf adaxial area (mm^2^; B, F, J), leaf dry matter content (mg/g; C, G, K), and leaf mass per area (mg/mm^2^; D, H, L) of the two sexes across the three dilution treatments (A, B, C, D), the four nitrogen treatments (E, F, G, H), and the four phosphorus treatments (I, J, K, L). Pairs of bars with different letters within a trait are significantly different from each other. Pairs of means with asterisks above are those in which females and hermaphrodites are significantly different from post-hoc comparisons of means.

## Discussion

Our aim in this study was to determine whether females and hermaphrodites differed in their growth responses to varying levels of overall nutrients, as well as relative levels of nitrogen and phosphorus. Using these data, we aimed to test for any cost of being a hermaphrodite in this system. We predicted that as nutrient availability decreases, both sexes should decrease in biomass, increase relative allocation to belowground growth, and the quality of their leaves in terms of nutrient concentrations should decrease. In addition, we predicted that hermaphrodite biomass, allocation, and leaf quality should change to a greater extent than those traits in females due to the costs associated with male function (pollen production and larger petals). We found that both sexes decreased biomass, increased relative allocation to belowground growth, and decreased leaf quality traits such as leaf nitrogen in response to decreased soil nutrients. However, these responses to decreased nutrients were not significantly greater for hermaphrodites. We discuss these findings further below.

### Lack of evidence for a biomass cost of male function

We did not find evidence that hermaphrodites decreased biomass or increased relative allocation to belowground growth more so than females in response to decreased nutrients. Male function in our species, therefore, does not appear to result in a significant biomass cost, at least prior to seed production, and at the nutrient levels used in this experiment. This finding contrasts with what has been shown in other species. When grown under nutrient-limiting conditions, females of *Plantago lanceolata* were larger than hermaphrodites overall, despite similar whole-plant nitrogen levels, suggesting that females used nitrogen that was not used for pollen production towards reproductive and vegetative biomass production (Poot, 1997). One reason for the differences between these two species could be the growth habit: *Plantago lanceolata* is a basal rosette perennial, while *Geranium maculatum* is a rhizomatous perennial with deciduous aboveground organs. It is possible that perennial species may have a lagged response to environmental nutrient changes due to the buffering effect of their perennial structures, such as rhizomes in our system. In addition, the *P*. *lanceolata* plants were each harvested 5 weeks after the onset of flowering for each individual plant, whereas our plants were harvested at the same time, when the first plant began to senesce. *Geranium maculatum* does not necessarily flower every year under all conditions, so we were unable to carry out a similar methodology. Regardless, it is important to note that the *P*. *lanceolata* plants were harvested at a later life history stage than those in this study, which could also account for the differences in results between the two experiments. Therefore, it is crucial to acknowledge that we were unable to directly measure whether the sexes differed in terms of demographic costs to reproduction, as we only measured plants after one season of reproduction, and not all plants produced flowers during this simulated season. However, biomass is a very common proxy for future reproduction in perennial plants (Younginger et al., 2017), so it is unlikely there would have been biologically meaningful differences in future years.

We also found that, at least at the leaf level, nitrogen uptake was similar between the sexes. This result is consistent with a study on another gynodioecious conger - *G*. *sylvaticum*. Asikainen & Mutikainen (2005) found that fertilizer additions increased seed production of both sex morphs equally in gynodioecious *Geranium sylvaticum*. In contrast, Eckhart and Chapin (1997) found that biomass of females in *Phacelia linearis* was only larger than that of hermaphrodites when grown in low nutrients, and that this difference disappeared when the sexes were grown at higher nutrient levels. It is possible that our lowest nutrient and lowest nitrogen levels may not have been low enough to detect differences between the sexes. More research on the nutrient relations of the two sexes in gynodioecious species would aid in our understanding of why some species show distinct differences in biomass across nutrient levels whereas others may not.

### Evidence for photosynthetic cost of male function

Contrary to the biomass results, we found evidence for a cost of male function in the response of leaf quality to nutrient levels. When growing under lower nitrogen treatments, both sexes had similar leaf chlorophyll contents. However, at higher relative nitrogen levels (most notably 8 mmol/L N), females had higher SPAD values than hermaphrodites, indicating that greater available nitrogen had benefited females more than hermaphrodites in terms of increasing leaf chlorophyll content. Assuming both sexes assimilated similar levels of soil nitrogen, this result suggests that the use of nitrogen for male function in hermaphrodites may come at a cost to leaf chlorophyll content. This finding is consistent with a study that more explicitly looked at photosynthetic rates of the two sexes in the gynodioecious species *Lobelia siphilitica*. Caruso et al. (2003) found that, in *L*. *siphilitica*, females had higher photosynthetic rates prior to reproduction than hermaphrodites. Chlorophyll content is expected to be positively correlated with maximum photosynthetic rates, at least at high light availability, so we might expect the same for our species at high light and high nitrogen environments (Buttery and Buzzell, 1977).

It is pertinent to highlight that, in a previous greenhouse experiment on *G*. *maculatum* in which photosynthetic rate was explicitly measured and plants were grown in higher nitrogen and high light conditions, there was no significant difference between the sexes in their photosynthetic rate (Van Etten et al., 2008). The data from this previous experiment does show a slight trend of higher photosynthetic rates in females than hermaphrodites under these conditions, so this discrepancy may be due to increased power in our current study (our degrees of freedom: 3, 46; theirs: 1, 11). It is possible that our incorporation of multiple populations, and higher sample size overall, allowed us to better able to detect sex differences under our study’s conditions. It is also relevant that in this experiment, we used sand, as opposed to a pine-bark vermiculite mix in the previous experiment. The distinct properties of these two soils may affect the mobility of nutrients, and thus could change how the plants respond. However, one would predict lower available nitrogen in the previous experiment due to the presence of clay particles, which can bind ammonium and affect the rate at which nitrogen is leached through the soil. Despite this fact, the measured per cent leaf nitrogen values in the previous experiment were on average higher than in this study, at a comparable life stage, which ultimately indicates that available soil nitrogen was in fact a little higher in their experiment. Furthermore, it is a distinct possibility that increased leaf SPAD values do not directly translate into higher photosynthetic rates in our species, as chlorophyll investment is only one part of realizing a high photosynthetic rate (Madrid et al., 2012). This could also explain the difference in pattern between these two studies, as again, the previous study directly measured photosynthetic rate. Further studies encompassing a more representative scale of genetic variation within the species and more explicit physiological measurements with controlled nutrient input could help elucidate the full extent of physiological differences and growth strategies of the two sexes in this species.

An important caveat of this study is that pollination did not occur in our experiment and hence we have no data for reproductive success. As *G*. *maculatum* is primarily an outcrossing species, we could not directly measure the relationship these different growth strategies may have on seed production in a greenhouse study. Our interpretation of the cost of male function (or lack thereof) should, thus, be viewed as a subset of the possible cost, excluding the potential effects on seed production – a commonly used framework for sex allocation theory.

## Conclusions

In summary, we found little evidence of a biomass cost of male function, despite evidence that hermaphrodites may have lower leaf chlorophyll content than females under high soil nitrogen conditions. The additional carbon gained from a greater photosynthetic capacity could be important for female seed production relative to that of hermaphrodites and could explain how females can compete against hermaphrodites in natural populations. Nevertheless, we only saw this difference at higher soil nitrogen levels. Hence this difference would only help explain female maintenance if sites with females and hermaphrodites are in fact not nitrogen-limited. In addition, these differences did not translate to biomass differences at this pre-seed production stage. The next step will be to investigate whether these differences in nutrient relations translate into fitness differences in specific ecological contexts, and how those patterns relate to variation in sex ratios in natural populations.

