## Supplemental Tables 1 and 2 for "Females and hermaphrodites of the gynodioecious *Geranium maculatum* respond similarly to soil nutrient availability"

**Supplemental Table 1**. Modified Hoagland's recipes (A) and the resulting moles of each element (B) used for N and P manipulation treatments. Each of these nitrogen and phosphorus treatments was either used as made (1x), or diluted to 0.5x or 0.1x of the original solution. Focal nutrient manipulations are highlighted with brackets.

**(A) Modified Hoagland's recipes**

| | Nitrogen treatments *(mL/L)* | | | Control *(mL/L)* | Phosphorus treatments *(mL/L)* | | |
|---|---|---|---|---|---|---|---|
| | **0.4 N** | **2 N** | **16 N** | **8 N ; 1 P** | **4 P** | **0.5 P** | **0.01 P** |
| **$KNO_3$** | [0.35] | [1] | [6] | 3 | 3 | 3 | 3 |
| **$Ca(NO_3)_2*4H_2O$** | [0] | [0.375] | [4] | 2 | 2 | 2 | 2 |
| **$K_2SO_4$** | [11.3] | [10] | [0] | 6 | [0] | [7] | [8] |
| **$KH_2PO_4$** | 2 | 2 | 2 | 2 | [8] | [1] | [0.02] |
| **$MgSO_4*7H_2O$** | 1 | 0 | 1 | 1 | 1 | 1 | 1 |
| **$CaSO_4*2H_2O$** | [400] | [1] | [0] | 200 | 200 | 200 | 200 |
| **NH4Cl** | [0.05] | [363] | [2] | 1 | 1 | 1 | 1 |
| **KCl**<br>**$H_3BO_3$**<br>**$MnSO_4*H_2O$**<br>**$ZnSO_4*7H_2O$**<br>**$CuSO_4*5H_2O$**<br>**MoO3**<br>**FeEDTA** | 0.5 | 0.5 | 0.5 | 0.5 | 0.5 | 0.5 | 0.5 |

**(B) Element moles in mmol/L**

| | **0.4 N** | **2 N** | **16 N** | **8 N ; 1 P** | **4 P** | **0.5 P** | **0.01 P** |
|---|---|---|---|---|---|---|---|
| **N** | [0.4] | [2] | [16] | 8 | 8 | 8 | 8 |
| **NO3-** | [0.35] | [1.75] | [14] | 7 | 7 | 7 | 7 |
| **NH4+** | [0.05] | [0.25] | [2] | 1 | 1 | 1 | 1 |
| **K** | 7 | 7 | 7 | 7 | 7 | 7 | 7.01 |
| **Ca** | 4 | 4.005 | 4 | 4 | 4 | 4 | 4 |
| **P** | 1 | 1 | 1 | 1 | [4] | [0.5] | [0.01] |
| **S** | 7.325 | 6.63 | 0.5 | 4 | 2.5 | 4.25 | 4.5 |
| **Mg** | 0.5 | 0.5 | 0.5 | 0.5 | 0.5 | 0.5 | 0.5 |
| **Cl** | 0.025 | 0.025 | 0.025 | 0.025 | 0.025 | 0.025 | 0.025 |
| **B** | 0.0125 | 0.0125 | 0.0125 | 0.0125 | 0.0125 | 0.0125 | 0.0125 |
| **Mn** | 0.001 | 0.001 | 0.001 | 0.001 | 0.001 | 0.001 | 0.001 |
| **Zn** | 0.001 | 0.001 | 0.001 | 0.001 | 0.001 | 0.001 | 0.001 |
| **Cu** | 0.00025 | 0.00025 | 0.00025 | 0.00025 | 0.00025 | 0.00025 | 0.00025 |
| **Mo** | 0.00025 | 0.00025 | 0.00025 | 0.00025 | 0.00025 | 0.00025 | 0.00025 |
| **Fe** | 0.025 | 0.025 | 0.025 | 0.025 | 0.025 | 0.025 | 0.025 |

**Supplemental Table 2a.** F-values, degrees of freedom, and p-values for the effects of the dilution, nitrogen, and phosphorus treatments, as well as their interactions with plant sex on biomass and other growth measurements. All models were run using a Gaussian error distribution.

| Data set | Dependent variable | Independent variables of interest | | | | | | | in. rh. mass |
|---|---|---|---|---|---|---|---|---|---|
| | | **Pop** | **Sex** | **N/P** | **Dilution** | **N/P*dil** | **Sex*dil** | **sex*N/P** | |
| all data | total dry mass estimate | F=54.91 | 4.31 | 1.51 | 4.85 | 2.40 | 1.96 | 1.93 | 411.70 |
| | | d.f=2, 138 | 1, 138 | 6, 138 | 2, 138 | 12, 138 | 2, 138 | 6, 138 | 1, 138 |
| | | **p<0.001** | **0.04** | **0.18** | **<0.01*** | **0.01** | **1.45** | **0.08** | **<0.001** |
| N man. | total dry mass estimate | 35.14 | 7.34 | 1.83 | 0.11 | 1.22 | 3.83 | 0.92 | 230.31 |
| | | 2, 79 | 1, 79 | 3, 79 | 2, 79 | 6, 79 | 2, 79 | 3, 79 | 1, 79 |
| | | **<0.001** | **0.01** | **0.15** | **0.89** | **0.30** | **0.03** | **0.43** | **<0.001** |
| P man. | total dry mass estimate | 26.20 | 0.05 | 0.20 | 8.76 | 2.06 | 1.15 | 2.13 | 175.07 |
| | | 2, 75 | 1, 75 | 3, 75 | 2, 75 | 6, 75 | 2, 75 | 3, 75 | 1, 75 |
| | | **<0.001** | **0.83** | **0.89** | **<0.001** | **0.07** | **0.32** | **0.10** | **<0.001** |
| all data | AG dry mass | 42.84 | 1.11 | 2.57 | 8.21 | 1.57 | 0.73 | 2.11 | 19.48 |
| | | 2, 109 | 1, 109 | 6, 109 | 2, 109 | 12, 109 | 2, 109 | 6, 109 | 1, 109 |
| | | **<0.001** | **0.29** | **0.02** | **<0.001*** | **0.11** | **0.48** | **0.06** | **<0.001** |
| N man. | AG dry mass | 24.94 | 0.13 | 5.42 | 3.36 | 1.01 | 1.34 | 1.50 | 4.29 |
| | | 2, 62 | 1, 62 | 3, 62 | 2, 62 | 6, 62 | 2, 62 | 3, 62 | 1, 62 |
| | | **<0.001** | **0.72** | **<0.01*** | **0.04** | **0.43** | **0.04** | **0.27** | **0.22** |
| P man. | AG dry mass | 23.08 | 3.65 | 0.09 | 7.96 | 0.-95 | 0.17 | 1.32 | 13.23 |
| | | 2, 60 | 1, 60 | 3, 60 | 2, 60 | 6, 60 | 2, 60 | 3, 60 | 1, 60 |
| | | **<0.001** | **0.06** | **0.96** | **<0.001** | **0.47** | **0.00** | **0.85** | **0.28** |
| all data | Final rhizome mass | 30.32 | 7.41 | 0.83 | 2.55 | 2.27 | 1.80 | 1.57 | 512.09 |
| | | 2, 138 | 1, 138 | 6, 138 | 2, 138 | 12, 138 | 2, 138 | 6, 138 | 1, 138 |
| | | **0.00** | **0.01** | **0.55** | **0.08** | **0.01** | **0.17** | **0.16** | **<0.001** |
| N man. | Final rhizome mass | 22.02 | 8.68 | 0.67 | 0.08 | 1.14 | 3.66 | 0.51 | 295.02 |
| | | 2, 79 | 1, 79 | 3, 79 | 2, 79 | 6, 79 | 2, 79 | 3, 79 | 1, 79 |
| | | **<0.001** | **0.00** | **0.57** | **0.92** | **0.35** | **0.03** | **0.68** | **<0.001** |
| P man. | Final rhizome mass | 13.90 | 0.89 | 0.49 | 6.42 | 2.29 | 1.00 | 2.67 | 232.16 |
| | | 2, 75 | 1, 75 | 3, 75 | 2, 75 | 6, 75 | 2, 75 | 3, 75 | 1, 75 |
| | | **<0.001** | **0.35** | **0.35** | **<0.01** | **0.04** | **0.37** | **0.05** | **<0.001** |
| all data | Highest recorded flower count (sq rt trans) | 28.80 | 0.77 | 1.87 | 1.96 | 1.56 | 1.14 | 1.61 | 8.94 |
| | | 2, 138 | 1, 138 | 6, 138 | 2, 138 | 12, 138 | 2, 138 | 6, 138 | 1, 138 |
| | | **<0.001** | **0.38** | **0.09** | **0.14** | **0.11** | **0.32** | **0.15** | **0.00** |
| N man. | Highest recorded flower count (sq rt trans) | 15.63 | 0.93 | 3.17 | 1.13 | 2.08 | 0.66 | 0.48 | 7.21 |
| | | 2, 79 | 1, 79 | 3, 79 | 2, 79 | 6, 79 | 2, 79 | 3, 79 | 1, 79 |
| | | **<0.001** | **0.34** | **0.03*** | **0.33** | **0.06** | **0.52** | **0.70** | **0.01** |
| P man. | Highest recorded flower count (sq rt trans) | 20.78 | 4.56 | 0.93 | 2.80 | 1.57 | 1.07 | 1.28 | 3.21 |
| | | 2, 75 | 1, 75 | 3, 75 | 2, 75 | 6, 75 | 2, 75 | 3, 75 | 1, 75 |
| | | **<0.001** | **0.04** | **0.43** | **0.07** | **0.17** | **0.35** | **0.29** | **0.08** |
| all data | Highest recorded leaf count (sq rt transformed) | 0.11 | 0.45 | 0.27 | 0.42 | 1.24 | 0.84 | 0.87 | 0.14 |
| | | 2, 138 | 1, 138 | 6, 138 | 2, 138 | 12, 138 | 2, 138 | 6, 138 | 1, 138 |
| | | **0.90** | **0.50** | **0.95** | **0.66** | **0.26** | **0.43** | **0.52** | **0.71** |
| N man. | Highest recorded leaf count (sq rt transformed) | 0.09 | 0.00 | 0.49 | 0.66 | 1.17 | 0.86 | 1.25 | 0.04 |
| | | 2, 79 | 1, 79 | 3, 79 | 2, 79 | 6, 79 | 2, 79 | 3, 79 | 1, 79 |
| | | **0.91** | **0.95** | **0.69** | **0.52** | **0.33** | **0.43** | **0.30** | **0.85** |
| P man. | Highest recorded leaf count (sq rt transformed) | 0.15 | 0.75 | 0.10 | 0.02 | 1.10 | 0.08 | 0.47 | 0.08 |
| | | 2, 75 | 1, 75 | 3, 75 | 2, 75 | 6, 75 | 2, 75 | 3, 75 | 1, 75 |
| | | **0.86** | **0.39** | **0.96** | **0.98** | **0.37** | **0.92** | **0.70** | **0.77** |

| **Data set** | **Dependent variable** | Independent variables of interest | | | | | | | **in. rh. mass** |
|---|---|---|---|---|---|---|---|---|---|
| | | **Pop** | **Sex** | **N/P** | **Dilution** | **N/P*dil** | **Sex*dil** | **sex*N/P** | |
| all data | BG : total dry mass | 56.62 | 0.85 | 2.37 | 3.84 | 0.82 | 1.94 | 1.20 | 6.48 |
| | | 2, 138 | 1, 138 | 6, 138 | 2, 138 | 12, 138 | 2, 138 | 6, 138 | 1, 138 |
| | | **<0.001** | **0.36** | **0.03** | **0.02*** | **0.63** | **0.15** | **0.31** | **0.01** |
| N man. | BG : total dry mass | 38.40 | 0.22 | 4.63 | 2.59 | 0.66 | 1.01 | 0.34 | 5.57 |
| | | 2, 79 | 1, 79 | 3, 79 | 2, 79 | 6, 79 | 2, 79 | 3, 79 | 1, 79 |
| | | **<0.001** | **0.64** | **<0.01*** | **0.08** | **0.69** | **0.37** | **0.80** | **0.02** |
| P man. | BG : total dry mass | 26.77 | 0.91 | 0.95 | 3.65 | 1.06 | 2.02 | 1.59 | 4.17 |
| | | 2, 75 | 1, 75 | 3, 75 | 2, 75 | 6, 75 | 2, 75 | 3, 75 | 1, 75 |
| | | **<0.001** | **0.34** | **0.42** | **0.03** | **0.39** | **0.04** | **0.14** | **0.20** |
| all data | infl : total dry mass (arcsin sq rt transformed) | 47.64 | 0.28 | 1.47 | 5.68 | 1.00 | 0.33 | 2.06 | 1.53 |
| | | 2, 138 | 1, 138 | 6, 138 | 2, 138 | 12, 138 | 2, 138 | 6, 138 | 1, 138 |
| | | **<0.001** | **0.60** | **0.19** | **<0.01*** | **0.45** | **0.72** | **0.06** | **0.22** |
| N man. | infl : total dry mass (arcsin sq rt transformed) | 26.87 | 0.83 | 2.44 | 3.81 | 1.24 | 0.48 | 2.22 | 0.92 |
| | | 2, 79 | 1, 79 | 3, 79 | 2, 79 | 6, 79 | 2, 79 | 3, 79 | 1, 79 |
| | | **<0.001** | **0.36** | **0.07** | **0.03** | **0.29** | **0.62** | **0.09** | **0.34** |
| P man. | infl : total dry mass (arcsin sq rt transformed) | 26.37 | 1.55 | 1.22 | 4.75 | 1.24 | 0.54 | 1.27 | 1.80 |
| | | 2, 75 | 1, 75 | 3, 75 | 2, 75 | 6, 75 | 2, 75 | 3, 75 | 1, 75 |
| | | **<0.001** | **0.22** | **0.31** | **0.01** | **0.29** | **0.18** | **0.58** | **0.29** |
| all data | inflorescence dry mass | 18.22 | 3.83 | 1.34 | 4.37 | 1.01 | 0.27 | 1.66 | 15.46 |
| | | 2, 105 | 1, 105 | 6, 105 | 2, 105 | 12, 105 | 2, 105 | 6, 105 | 1, 105 |
| | | **<0.001** | **0.05** | **0.24** | **0.02*** | **0.45** | **0.00** | **0.76** | **0.14** |
| N man. | inflorescence dry mass | 11.13 | 0.14 | 3.34 | 3.24 | 1.52 | 0.44 | 2.24 | 6.31 |
| | | 2, 58 | 1, 58 | 3, 58 | 2, 58 | 6, 58 | 2, 58 | 3, 58 | 1, 58 |
| | | **<0.001** | **0.71** | **0.03*** | **0.05** | **0.19** | **0.65** | **0.09** | **0.01** |
| P man. | inflorescence dry mass | 10.20 | 7.33 | 0.19 | 4.00 | 0.75 | 1.29 | 0.41 | 14.08 |
| | | 2, 59 | 1, 59 | 3, 59 | 2, 59 | 6, 59 | 2, 59 | 3, 59 | 1, 59 |
| | | **0.00** | **0.01** | **0.91** | **0.02** | **0.61** | **0.00** | **0.28** | **0.75** |
| all data | leaf area | 8.74 | 0.09 | 1.03 | 1.13 | 1.63 | 0.20 | 0.57 | 2.05 |
| | | 2, 74 | 1, 74 | 6, 74 | 2, 74 | 12, 74 | 2, 74 | 6, 74 | 1, 74 |
| | | **0.00** | **0.76** | **0.41** | **0.33** | **0.10** | **0.82** | **0.75** | **0.16** |
| N man. | leaf area | 4.05 | 0.62 | 1.87 | 0.12 | 1.58 | 1.94 | 1.10 | 0.49 |
| | | 2, 41 | 1, 41 | 3, 41 | 2, 41 | 6, 41 | 2, 41 | 3, 41 | 1, 41 |
| | | **0.02** | **0.44** | **0.15** | **0.89** | **0.18** | **0.49** | **0.16** | **0.36** |
| P man. | leaf area | 6.97 | 0.01 | 0.36 | 3.84 | 1.15 | 1.10 | 0.18 | 1.12 |
| | | 2, 36 | 1, 36 | 3, 36 | 2, 36 | 6, 36 | 2, 36 | 3, 36 | 1, 36 |
| | | **0.00** | **0.93** | **0.78** | **0.03** | **0.36** | **0.34** | **0.91** | **0.30** |
| all data | leaf dry matter content | 3.69 | 0.46 | 1.49 | 0.25 | 0.87 | 0.47 | 0.56 | 1.25 |
| | | 2, 76 | 1, 76 | 6, 76 | 2, 76 | 12, 76 | 2, 76 | 6, 76 | 1, 76 |
| | | **0.03** | **0.50** | **0.19** | **0.78** | **0.58** | **0.27** | **0.63** | **0.76** |
| N man. | leaf dry matter content | 2.63 | 1.01 | 1.71 | 0.34 | 1.05 | 0.49 | 0.34 | 0.42 |
| | | 2, 41 | 1, 41 | 3, 41 | 2, 41 | 6, 41 | 2, 41 | 3, 41 | 1, 41 |
| | | **0.08** | **0.32** | **0.18** | **0.72** | **0.41** | **0.61** | **0.80** | **0.52** |
| P man. | leaf dry matter content | 4.25 | 0.58 | 0.83 | 0.75 | 0.55 | 0.26 | 0.54 | 3.78 |
| | | 2, 39 | 1, 39 | 3, 39 | 2, 39 | 6, 39 | 2, 39 | 3, 39 | 1, 39 |
| | | **0.02** | **0.45** | **0.48** | **0.48** | **0.77** | **0.78** | **0.66** | **0.06** |

"N man." = data for when nitrogen was manipulated; "P man."= data for when phosphorus was manipulated; "error dist."= error distribution used. "in. rh. mass" = initial rhizome mass; "N/P"= the effect of nitrogen manipulation, phosphorus manipulation, or both, dependent on the data set; "Pop"= plant source population; *=p-value was significant after Benjamini-Hochberg test.

**Supplemental Table 2b.** F-values, degrees of freedom, and p-values for the effects of the dilution, nitrogen, and phosphorus treatments, as well as their interactions with plant sex on biomass, allocation, and leaf quality measurements. Values in black text indicate effects of interest for a given model.

| **Data set** | **Dependent variable** | Independent variables of interest | | | | | | | **in. rh. mass** |
|---|---|---|---|---|---|---|---|---|---|
| | | **Pop** | **Sex** | **N/P** | **Dilution** | **N/P*dil** | **Sex*dil** | **sex*N/P** | |
| all data | BG : total dry mass | F=56.62 | 0.85 | 2.37 | 3.84 | 0.82 | 1.94 | 1.20 | 6.48 |
| | | d.f=2, 138 | 1, 138 | 6, 138 | 2, 138 | 12, 138 | 2, 138 | 6, 138 | 1, 138 |
| | | **p<0.001** | **0.36** | **0.03** | **0.02*** | **0.63** | **0.15** | **0.31** | **0.01** |
| N man. | BG : total dry mass | 38.40 | 0.22 | 4.63 | 2.59 | 0.66 | 1.01 | 0.34 | 5.57 |
| | | 2, 79 | 1, 79 | 3, 79 | 2, 79 | 6, 79 | 2, 79 | 3, 79 | 1, 79 |
| | | **<0.001** | **0.64** | **<0.01*** | **0.08** | **0.69** | **0.37** | **0.80** | **0.02** |
| P man. | BG : total dry mass | 26.77 | 0.91 | 0.95 | 3.65 | 1.06 | 2.02 | 1.59 | 4.17 |
| | | 2, 75 | 1, 75 | 3, 75 | 2, 75 | 6, 75 | 2, 75 | 3, 75 | 1, 75 |
| | | **<0.001** | **0.34** | **0.42** | **0.03** | **0.39** | **0.04** | **0.14** | **0.20** |
| all data | infl : total dry mass (arcsin sq rt transformed) | 47.64 | 0.28 | 1.47 | 5.68 | 1.00 | 0.33 | 2.06 | 1.53 |
| | | 2, 138 | 1, 138 | 6, 138 | 2, 138 | 12, 138 | 2, 138 | 6, 138 | 1, 138 |
| | | **<0.001** | **0.60** | **0.19** | **<0.01*** | **0.45** | **0.72** | **0.06** | **0.22** |
| N man. | infl : total dry mass (arcsin sq rt transformed) | 26.87 | 0.83 | 2.44 | 3.81 | 1.24 | 0.48 | 2.22 | 0.92 |
| | | 2, 79 | 1, 79 | 3, 79 | 2, 79 | 6, 79 | 2, 79 | 3, 79 | 1, 79 |
| | | **<0.001** | **0.36** | **0.07** | **0.03** | **0.29** | **0.62** | **0.09** | **0.34** |
| P man. | infl : total dry mass (arcsin sq rt transformed) | 26.37 | 1.55 | 1.22 | 4.75 | 1.24 | 0.54 | 1.27 | 1.80 |
| | | 2, 75 | 1, 75 | 3, 75 | 2, 75 | 6, 75 | 2, 75 | 3, 75 | 1, 75 |
| | | **<0.001** | **0.22** | **0.31** | **0.01** | **0.29** | **0.18** | **0.58** | **0.29** |
| all data | inflorescence dry mass | 18.22 | 3.83 | 1.34 | 4.37 | 1.01 | 0.27 | 1.66 | 15.46 |
| | | 2, 105 | 1, 105 | 6, 105 | 2, 105 | 12, 105 | 2, 105 | 6, 105 | 1, 105 |
| | | **<0.001** | **0.05** | **0.24** | **0.02*** | **0.45** | **0.76** | **0.14** | **0.002** |
| N man. | inflorescence dry mass | 11.13 | 0.14 | 3.34 | 3.24 | 1.52 | 0.44 | 2.24 | 6.31 |
| | | 2, 58 | 1, 58 | 3, 58 | 2, 58 | 6, 58 | 2, 58 | 3, 58 | 1, 58 |
| | | **<0.001** | **0.71** | **0.03*** | **0.05** | **0.19** | **0.65** | **0.09** | **0.01** |
| P man. | inflorescence dry mass | 10.20 | 7.33 | 0.19 | 4.00 | 0.75 | 1.29 | 0.41 | 14.08 |
| | | 2, 59 | 1, 59 | 3, 59 | 2, 59 | 6, 59 | 2, 59 | 3, 59 | 1, 59 |
| | | **0.00** | **0.01** | **0.91** | **0.02** | **0.61** | **0.00** | **0.28** | **0.75** |
| all data | leaf area | 8.74 | 0.09 | 1.03 | 1.13 | 1.63 | 0.20 | 0.57 | 2.05 |
| | | 2, 74 | 1, 74 | 6, 74 | 2, 74 | 12, 74 | 2, 74 | 6, 74 | 1, 74 |
| | | **0.00** | **0.76** | **0.41** | **0.33** | **0.10** | **0.82** | **0.75** | **0.16** |
| N man. | leaf area | 4.05 | 0.62 | 1.87 | 0.12 | 1.58 | 1.94 | 1.10 | 0.49 |
| | | 2, 41 | 1, 41 | 3, 41 | 2, 41 | 6, 41 | 2, 41 | 3, 41 | 1, 41 |
| | | **0.02** | **0.44** | **0.15** | **0.89** | **0.18** | **0.49** | **0.16** | **0.36** |
| P man. | leaf area | 6.97 | 0.01 | 0.36 | 3.84 | 1.15 | 1.10 | 0.18 | 1.12 |
| | | 2, 36 | 1, 36 | 3, 36 | 2, 36 | 6, 36 | 2, 36 | 3, 36 | 1, 36 |
| | | **0.00** | **0.93** | **0.78** | **0.03** | **0.36** | **0.34** | **0.91** | **0.30** |
| all data | leaf dry matter content | 3.69 | 0.46 | 1.49 | 0.25 | 0.87 | 0.47 | 0.56 | 1.25 |
| | | 2, 76 | 1, 76 | 6, 76 | 2, 76 | 12, 76 | 2, 76 | 6, 76 | 1, 76 |
| | | **0.03** | **0.50** | **0.19** | **0.78** | **0.58** | **0.27** | **0.63** | **0.76** |
| N man. | leaf dry matter content | 2.63 | 1.01 | 1.71 | 0.34 | 1.05 | 0.49 | 0.34 | 0.42 |
| | | 2, 41 | 1, 41 | 3, 41 | 2, 41 | 6, 41 | 2, 41 | 3, 41 | 1, 41 |
| | | **0.08** | **0.32** | **0.18** | **0.72** | **0.41** | **0.61** | **0.80** | **0.52** |
| P man. | leaf dry matter content | 4.25 | 0.58 | 0.83 | 0.75 | 0.55 | 0.26 | 0.54 | 3.78 |
| | | 2, 39 | 1, 39 | 3, 39 | 2, 39 | 6, 39 | 2, 39 | 3, 39 | 1, 39 |
| | | **0.02** | **0.45** | **0.48** | **0.48** | **0.77** | **0.78** | **0.66** | **0.06** |

"N man." = data for when nitrogen was manipulated; "P man."= data for when phosphorus was manipulated; "error dist."= error distribution used. "in. rh. mass" = initial rhizome mass; "N/P"= the effect of nitrogen manipulation, phosphorus manipulation, or both, dependent on the data set; "Pop"= plant source population; *=p-value was significant after Benjamini-Hochberg test.

**Supplemental Table 2c.** F-values, degrees of freedom, and p-values for the effects of the dilution, nitrogen, and phosphorus treatments, as well as their interactions with plant sex on leaf quality measurements. Values in black text indicate effects of interest for a given model.

| **Data set** | **Dependent variable** | Independent variables of interest | | | | | | | **in. rh. mass** |
|---|---|---|---|---|---|---|---|---|---|
| | | **Pop** | **Sex** | **N/P** | **Dilution** | **N/P*dil** | **Sex*dil** | **sex*N/P** | |
| all data | LMA | F=1.90 | 1.21 | 0.97 | 0.51 | 0.67 | 0.56 | 0.74 | 0.65 |
| | | d.f.=2, 69 | 1, 69 | 6, 69 | 2, 69 | 12, 69 | 2, 69 | 6, 69 | 1, 69 |
| | | **p=0.16** | **0.28** | **0.45** | **0.61** | **0.77** | **0.57** | **0.62** | **0.42** |
| N man. | LMA | 2.03 | 1.30 | 1.14 | 0.24 | 0.87 | 0.66 | 0.60 | 0.33 |
| | | 2, 38 | 1, 38 | 3, 38 | 2, 38 | 6, 38 | 2, 38 | 3, 38 | 1, 38 |
| | | **0.15** | **0.26** | **0.35** | **0.78** | **0.52** | **0.52** | **0.62** | **0.57** |
| P man. | LMA | 1.19 | 0.14 | 0.65 | 0.17 | 0.43 | 0.08 | 1.06 | 2.20 |
| | | 2, 34 | 1, 34 | 3, 34 | 2, 34 | 6, 34 | 2, 34 | 3, 34 | 1, 34 |
| | | **0.32** | **0.71** | **0.59** | **0.84** | **0.85** | **0.15** | **0.93** | **0.38** |
| all data | % leaf N | 3.45 | 0.00 | 3.40 | 5.07 | 1.74 | 0.60 | 0.90 | 0.01 |
| | | 2, 64 | 1, 64 | 6, 64 | 2, 64 | 12, 64 | 2, 64 | 6, 64 | 1, 64 |
| | | **0.04** | **0.98** | **0.01** | **<0.01*** | **0.08** | **0.55** | **0.50** | **0.93** |
| N man. | % leaf N | 5.29 | 1.33 | 4.84 | 1.23 | 0.65 | 0.93 | 0.15 | 0.05 |
| | | 2, 37 | 1, 37 | 3, 37 | 2, 37 | 6, 37 | 2, 37 | 3, 37 | 1, 37 |
| | | **0.01** | **0.26** | **<0.01*** | **0.30** | **0.69** | **0.40** | **0.93** | **0.83** |
| P man. | % leaf N | 0.10 | 0.72 | 1.98 | 7.05 | 1.76 | 0.08 | 2.15 | 0.07 |
| | | 2, 31 | 1, 31 | 3, 31 | 2, 31 | 6, 31 | 2, 31 | 3, 31 | 1, 31 |
| | | **0.91** | **0.40** | **0.14** | **<0.001** | **0.14** | **0.93** | **0.11** | **0.80** |
| all data | % leaf P | 1.61 | 0.32 | 1.29 | 1.17 | 1.02 | 1.13 | 1.23 | 1.75 |
| | | 2, 39 | 1, 39 | 6, 39 | 2, 39 | 12, 39 | 2, 39 | 6, 39 | 1, 39 |
| | | **0.21** | **0.57** | **0.28** | **0.32** | **0.45** | **0.33** | **0.31** | **0.19** |
| N man. | % leaf P | 1.93 | 0.21 | 0.58 | 2.78 | 0.61 | 0.48 | 0.62 | 3.22 |
| | | 2, 21 | 1, 21 | 3, 21 | 2, 21 | 6, 21 | 2, 21 | 3, 21 | 1, 21 |
| | | **0.17** | **0.65** | **0.64** | **0.08** | **0.72** | **0.63** | **0.61** | **0.09** |
| P man. | % leaf P | 0.11 | 0.05 | 1.78 | 0.65 | 0.81 | 0.31 | 1.82 | 0.16 |
| | | 2, 20 | 1, 20 | 3, 20 | 2, 20 | 6, 20 | 2, 20 | 3, 20 | 1, 20 |
| | | **0.90** | **0.83** | **0.18** | **0.53** | **0.58** | **0.69** | **0.88** | **0.18** |
| all data | Leaf N:P ratio | 3.43 | 3.14 | 1.16 | 0.09 | 1.31 | 0.77 | 0.98 | 2.21 |
| | | 2, 36 | 1, 36 | 6, 36 | 2, 36 | 11, 36 | 2, 36 | 5, 36 | 1, 36 |
| | | **0.04** | **0.09** | **0.35** | **0.91** | **0.26** | **0.15** | **0.47** | **0.44** |
| N man. | Leaf N:P ratio | 1.83 | 1.26 | 0.87 | 0.50 | 0.74 | 0.75 | 0.60 | 2.03 |
| | | 2, 19 | 1, 19 | 3, 19 | 2, 19 | 6, 19 | 2, 19 | 3, 19 | 1, 19 |
| | | **0.19** | **0.28** | **0.47** | **0.61** | **0.62** | **0.49** | **0.62** | **0.17** |
| P man. | Leaf N:P ratio | 0.41 | 2.01 | 1.38 | 0.55 | 0.72 | 0.30 | 1.21 | 0.92 |
| | | 2, 19 | 1, 19 | 3, 19 | 2, 19 | 5, 19 | 2, 19 | 2, 19 | 1, 19 |
| | | **0.67** | **0.17** | **0.28** | **0.58** | **0.61** | **0.35** | **0.74** | **0.32** |
| all data | SPAD | 2.42 | 2.49 | 1.49 | 0.22 | 1.58 | 0.29 | 2.37 | 0.00 |
| | | 2, 80 | 1, 80 | 6, 80 | 2, 80 | 12, 80 | 2, 80 | 6, 80 | 1, 80 |
| | | **0.10** | **0.12** | **0.19** | **0.81** | **0.12** | **0.75** | **0.04** | **0.97** |
| N man. | SPAD | 0.30 | 4.75 | 4.08 | 0.98 | 3.36 | 1.73 | 5.81 | 0.08 |
| | | 2, 46 | 1, 46 | 3, 46 | 2, 46 | 6, 46 | 2, 46 | 3, 46 | 1, 46 |
| | | **0.74** | **0.03** | **0.01*** | **0.38** | **0.01** | **0.19** | **<0.01*** | **0.78** |
| P man. | SPAD | 2.59 | 1.74 | 0.97 | 0.89 | 0.33 | 0.69 | 1.12 | 0.28 |
| | | 2, 39 | 1, 39 | 3, 39 | 2, 39 | 6, 39 | 2, 39 | 3, 39 | 1, 39 |
| | | **0.09** | **0.19** | **0.42** | **0.42** | **0.91** | **0.51** | **0.35** | **0.60** |

"N man." = data for when nitrogen was manipulated; "P man."= data for when phosphorus was manipulated; "error dist."= error distribution used. "in. rh. mass" = initial rhizome mass; "N/P"= the effect of nitrogen manipulation, phosphorus manipulation, or both, dependent on the data set; "Pop"= plant source population; *=p-value was significant after Benjamini-Hochberg test.
